# Beyond a diagnostic tool: Validating standardized Mahalanobis distance as a species distribution model for invasive alien species in North America

**DOI:** 10.1101/2024.04.30.591852

**Authors:** JR Stinziano, A Charron, M Damus

**Affiliations:** Plant Health Science Directorate, Canadian Food Inspection Agency; Ottawa Plant Laboratory – Genotyping and Botany, Canadian Food Inspection Agency

## Abstract

Species distribution models (SDMs) are useful tools for predicting where new invasive species can establish within a country, supporting both preparatory and response activities. National Plant Protection Organizations use SDMs to inform risk assessment and surveillance activities for emerging plant pests. However, SDMs face multiple and difficult statistical challenges, including multi-collinearity of input variables, correlation structures in climate variables that vary through time and space, limited species observation data, and they often lack formal tests of model performance.

We have implemented a previously-reported extrapolation-detection tool as an SDM, rather than a diagnostic tool of SDMs. This method characterizes the observed multivariate climate envelope by using Mahalanobis distance to take advantage of the correlation between climate variables, and identifies areas where the climatic conditions are outside the range of the observed climate envelope. Model outputs include climate suitability maps, and most-important covariate analyses to identify the environmental drivers of the results while assisting in variable reduction. We performed a formal test to assess the ability of the SDM to identify areas invaded by invasive plant pests in North America.

Using a list of 23 species from the Canadian Food Inspection Agency’s regulated plant pest list, we demonstrate that this method achieves a high level of accuracy (> 85%) for determining climate suitability for North American plant pest invasions, especially when combined with most-important covariate-guided variable reduction. This suggests that the model is suitable for identifying areas of North America that are susceptible to future invasions. We show that many of the errors occur at the edge of climate suitable areas, where we would expect greater uncertainty in model predictions due to potential over-constraining and geospatial averaging. We present additional analyses to support recommendations on the use and limitations of this SDM in a regulatory context.

## Introduction

Species distribution models are an important component of the invasive pest management toolkit (Srivastava et al., 2019). These models provide a means to identify where a pest may survive, which can be used to inform risk assessment, monitoring, containment, and eradication activities as well as help National Plant Protection Organizations make regulatory decisions (e.g. Kendig et al., 2022; Venette, 2015).

There are two primary categories of SDMs: process-based and correlative (Srivastava et al., 2019). Process-based models (e.g. CLIMEX “Match locations”, Sutherst and Maywald, 1985, Sutherst et al., 2007) use biologically-derived parameters and climate data along with species presence data to predict species distributions. Correlative models (e.g. Climex “Match climates” and “Regional match climates”, the Climatch algorithm, Crombie et al., 2008, Kriticos et al., 2015; MAXENT, e.g. Elith et al., 2010; and Self-Organizing Maps, e.g. Worner and Gevrey, 2006) use the climate and species presence data alone to predict areas of climate suitability for the species. Performance of both process-based and correlative models can vary depending on input datasets (Ahmadi et al., 2023; Early et al., 2022). However, correlative model algorithms require less data and often less detailed or experimentally-derived biological knowledge of a species than process-based models, and have advantages of implementation speed at the cost of potentially reduced biological realism, projection uncertainty (e.g. Webber et al., 2011, Jarnevich et al., 2015 and Jarnevich & Young, 2015) and issues with collinearity within the climate data (e.g. Dormann et al., 2013). Rapid implementation and lower data requirements can favour the use of correlative over process-based models for emerging pest species where biological information may be scarce and the timeframe to react may be short.

Mesgaran et al. (2014) developed a method to assess extrapolation in correlative SDMs, which can identify univariate novelty and novel combinations of covariates. Univariate novelty occurs where the SDM predicts climatic suitability outside of the range of at least one climatic variable (e.g. predicting climate suitability where the temperature is hotter than the observed climate envelope for the species). Novel combinations of covariates occur when the SDM predicts climatic suitability in a location where the climate is within univariate tolerances of the observed climate envelope, but has a combination of climate variables that doesn’t exist within the observed climate envelope (e.g. predicting climate suitability in an area that is ‘hot’ and ‘wet’ because the species has been observed in hot/dry and cold/wet environments, but not specifically a hot/wet environment). Mesgaran et al. (2014) present their method as a diagnostic tool for correlative SDM projections.

Here we propose using the Mesgaran et al. (2014) method as a correlative SDM. Our purpose is to assess the validity of using the standardized Mahalanobis distances for identifying areas of potential invasion for plant pest species. Our objectives were to:

1. Test the method on existing invasions so that the method can be used to predict potential invasion sites in the future.
2. Provide variability metrics for climate suitability that are useful in supporting pest risk assessments and pest management decisions.
3. Assess limitations on the SDM due to characteristics of the input dataset.

## Material and Methods

### Climate Data

The CHELSA 1981-2010 climate dataset version 2.1 (Beck et al., 2020; Brun et al. preprint; Brun et al., 2022; Karger et al. 2017; Karger & Zimmermann, 2018a; Karger et al., 2018b; Karger et al., 2019; Karger et al., 2020a; Karger et al., 2020b) was used. We used two different sets of bioclimatic variables as determined by plant health risk assessors at the Canadian Food Inspection Agency, based on whether the pest was an insect or plant (Table 1). Raster data was downscaled from 30 arc-second resolution to 10 arc-minute resolution.

**Table 1.** – List of bioclimatic variables used for species distribution modeling by organism type.

| Organism Type | Temperature-related Bioclimatic Variables | Precipitation-related Bioclimatic Variables |
| --- | --- | --- |
| Insect | bio1, bio5, bio6, bio8, bio9, bio10, bio11 | bio12, bio13, bio14, bio16, bio17, bio18, bio19 |
| Plant | bio1, bio2, bio4, bio8, bio9, bio10, bio11 | bio12, bio13, bio14, bio15, bio18, bio19 |

bio1: mean annual air temperature; bio2: mean diurnal air temperature range; bio4: temperature seasonality; bio5: mean daily maximum air temperature of the warmest month; bio6: mean daily minimum air temperature of the coldest month; bio8: mean daily air temperatures of the wettest quarter; bio9: mean daily air temperatures of the driest quarter; bio10: mean daily air temperatures of the warmest quarter; bio11: mean daily air temperatures of the coldest quarter; bio12: annual precipitation amount; bio13: precipitation amount of the wettest month; bio14: precipitation amount of the driest month; bio15: precipitation seasonality; bio16: mean monthly precipitation of the wettest quarter; bio17: mean monthly precipitation of the driest quarter; bio18: mean monthly precipitation of the warmest quarter; bio19: mean monthly precipitation of the coldest quarter (Karger et al., 2017).

The range of values for each species across the bioclimatic variables for the reference climate are available in Table S 1 and Table S 2.

### Species Occurrence Data

We targeted species on the Canadian Food Inspection Agency’s regulated pest list (Table 2). Occurrence data were obtained from GBIF (GBIF.org) using rgbif (Chamberlain et al., 2023; Chamberlain and Boettiger, 2017). Occurrence data was then aggregated to 10 arc-minute pixels. Occurrence data was then partitioned into a training dataset of all observations outside North America, and a testing dataset of all observations within North America to support model evaluation as per Sofaer et al. (2019).

**Table 2.** – Species selected for model validation, including distribution of observations between training (non-North American observations) and test (North American observation) datasets.

| Species | Number of Observations |  | Number of Pixels with Observations |  | GBIF Data DOI(s) |
| --- | --- | --- | --- | --- | --- |
|  | Training | Test | Training | Test |  |
| *Aegilops cylindrica* | 2,142 | 318 | 653 | 180 | 10.15468/dl.sxf4ed;<br>10.15468/dl.apr26s |
| *Alopecurus myosuroides* | 77,926 | 9 | 3,694 | 2 | 10.15468/dl.yrms5h;<br>10.15468/dl.kj8avq;<br>10.15468/dl.rqmx8s |
| *Arundo donax* | 34,157 | 4,396 | 2,604 | 1,338 | 10.15468/dl.6sewx4 |
| *Centaurea iberica* | 615 | 32 | 159 | 19 | 10.15468/dl.6sewx4 |
| *Centaurea solstitialis* | 8,594 | 3,524 | 2,095 | 630 | 10.15468/dl.6sewx4 |
| *Crupina vulgaris* | 9,373 | 82 | 1,052 | 26 | 10.15468/dl.6sewx4 |
| *Echium plantagineum* | 58,597 | 124 | 5,335 | 31 | 10.15468/dl.6sewx4 |
| *Eriochloa villosa* | 559 | 151 | 299 | 79 | 10.15468/dl.bjarjw |
| *Microstegium vimineum* | 1,958 | 10,568 | 584 | 1,953 | 10.15468/dl.z3vgwm |
| *Paspalum dilatatum* | 32,473 | 2,072 | 3,016 | 873 | 10.15468/dl.6sewx4 |
| *Persicaria perfoliata* | 660 | 4,556 | 271 | 472 | 10.15468/dl.6sewx4 |
| *Pueraria montana* | 4,465 | 4,760 | 895 | 1240 | 10.15468/dl.xjskcb;<br>10.15468/dl.fv3kyq;<br>10.15468/dl.m64wtj;<br>10.15468/dl.x32d76 |
| *Senecio inaequidens* | 330,860 | 793 | 3,593 | 142 | 10.15468/dl.6sewx4 |
| *Zygophyllum fabago* | 1,566 | 36 | 296 | 18 | 10.15468/dl.6sewx4 |
| *Cacoecimorpha pronubana* | 7,974 | 8 | 729 | 7 | 10.15468/dl.6sewx4 |
| *Cydalima perspectalis* | 153,043 | 270 | 3,664 | 40 | 10.15468/dl.sc8naw |
| *Epiphyas postvittana* | 34,482 | 144 | 608 | 38 | 10.15468/dl.6sewx4 |
| *Lobesia botrana* | 532 | 7 | 238 | 7 | 10.15468/dl.6sewx4 |
| *Lycorma delicatula* | 1,245 | 16,860 | 736 | 399 | 10.15468/dl.tfq3yz |
| *Popillia japonica* | 5,308 | 23,623 | 176 | 3,875 | 10.15468/dl.6sewx4 |
| *Rhagoletis cerasi* | 730 | 46 | 354 | 23 | 10.15468/dl.tkebwz;<br>10.15468/dl.hsqxzv |
| *Sirex noctilio* | 240 | 532 | 172 | 19 | 10.15468/dl.6sewx4 |
| *Tetropium fuscum* | 1,032 | 327 | 427 | 75 | 10.15468/dl.b3ne3v;<br>10.15468/dl.ny7zwh |

Canadian Food Inspection Agency laboratory diagnostics and survey data were included in the test of the model for species that have been detected in Canada, where the observations were not in nurseries, and where the detection was not associated directly with the movement of goods into Canada (i.e. during import commodity inspections).

To be included in the analysis, species had to meet the following criteria:

1. Be on the Canadian Food Inspection Agency’s list of regulated pests (CFIA, 2024).
2. Have observations in raster pixels totaling at least 10 X the number of bioclimatic variables within the training dataset (i.e. observations in at least 140 pixels for invertebrates, at least 130 pixels for plants) (as per the recommendations of Jarnevich et al., 2015).
3. Have observations in North America to validate the model.

### Mahalanobis distance, Type 1, and Type 2 Novelty

Mesgaran et al. (2014) proposed a method to detect extrapolation in SDMs. First, the univariate distances of points in the SDM projection from the reference climate for each climate variable are calculated (See Table S 1, Table S 2, and Table S 3 for the reference climate for each species), and then summed to determine Type 1 Novelty:

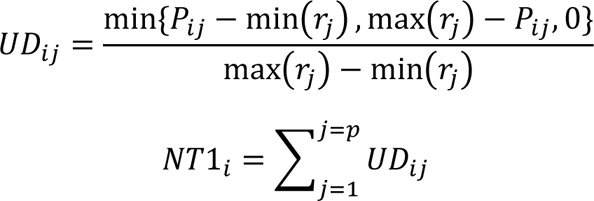

Where *UD_ij_* is the univariate distance from projection grid cell *i* for covariate *j*, *P_ij_* is the value of grid cell *i* for covariate *j* in the projection data, *min(r_j_)* and *max(r_j_)* are the minimum and maximum values of covariate *j* for the reference data *r*, and *NT1_i_* is the type 1 novelty value for grid cell *i*. *NT1* falls within the domain of negative infinity and 0, with values of 0 indicating grid cells within the univariate range of the climatic data, and all negative values indicating that the grid cell falls outside the univariate range.

For all grid cells where *NT1* = 0, type 2 novelty (*NT2*) is calculated using Mahalanobis distances. Mahalanobis distance can be used to detect outliers in a multivariate space, and can be calculated as:

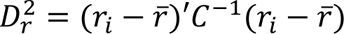

Where *D_r_^2^* is Mahalanobis distance of the data in the reference data, and *‘C^-1^* is the covariance matrix. Type 2 Novelty (*NT2*) can then be calculated as:

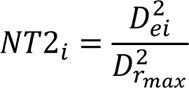

Where *D^2^_rmax_* is the maximum Mahalanobis distance from the reference dataset, and *D^2^_ei_* is the Mahalanobis distance of point *e_i_* from *r̅*, calculated as:

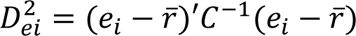

*NT2* values vary from 0 to positive infinity, with values between 0 and 1 being considered climate suitable.

### Most Important Variable Calculations

The most important variable calculation for each pixel was run according to Mesgaran et al. (2014), and used to construct a categorical raster. Briefly, covariate importance was calculated as:

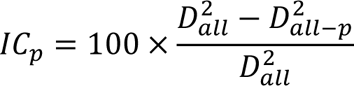

Where *IC_p_* is the importance of covariate *p* at a point, *D^2^_all_* is the Mahalanobis distance with all covariates at a given point, and *D^2^_all-p_* is the Mahalanobis distance with all covariates except for covariate p at a given point. The most important covariate is then the covariate with the highest *IC_p_* value at a given point.

### Invasion Performance Testing

For invasion performance testing, the testing dataset for species occurrence data was rasterized into a binary 1/0 raster. This was then used to filter the ensemble predictions from the training data to categorize the test data into climate suitable, type 1 novelty, or type 2 novelty. Multivariate accuracy was calculated based on pixels within the test dataset as:

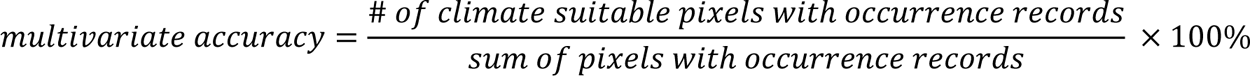

False negative rate was calculated from the test dataset as:

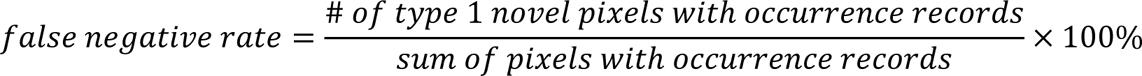

Finally, univariate accuracy was calculated as:

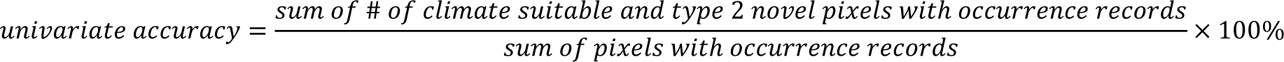

### Stepwise Most Important Covariate (MIC) Analysis

For species where multivariate accuracy was low, we performed a stepwise reduction in climate variables, dropping the most important covariate that led to Type 2 Novelty estimates in areas where the species is found. This follows the logic that if a prediction is incorrect, the most important covariate is a likely limitation on the accuracy of the prediction. We performed this stepwise process until multivariate accuracy exceeded 85%.

### Invasion Hotspot Map

An invasion hotspot map was created by converting the climate suitability rasters into binary rasters, with 1 representing multivariate climate suitability, and 0 representing both type 1 and type 2 novel conditions. The binary raster layers for all species were then added together to identify hotspots where multiple species had multivariate climate suitability.

### iNaturalist Plant Data Review

A randomized subset of the “research-grade” data obtained through GBIF from iNaturalist from the training datasets was spot-checked by a trained botanist (AC), to verify correct species identifications for *Eriochloa villosa* and *Alopecurus myosuroides* based on photo identification. To properly identify these species, a clear view of the inconspicuous flowers and/or surrounding characteristics are necessary and as such is dependent on the user’s knowledge about the important characteristics to capture and on photograph quality (e.g. angle, sharpness and distance). The species *Aegilops cylindrica*, *Microstegium vimineum* and *Pueraria montana* have sufficient conspicuous characteristics to allow proper identification and are usually captured in photographs due to their salient traits (e.g. flowers, fruits and growing nature of *P. montana*). As highlighted in White et al. (2023), careful consideration is essential when dealing with species identifications that depend on intricate morphological details (e.g. Cyperaceae, Poaceae, bryophytes). The objective of this exercise was to account for the potential error present in the model. As such, only *E. villosa* and *A. myosuroides* were assessed. The observations were classified as “accurate” if the observation was taxonomically correct, “inaccurate” if the observation was clearly misidentified and “ambiguous” if, in our opinion, the observation could not have been made with the available information (e.g. photograph taken from too far away, too blurry or taxonomically distinct characteristics not included or blurry). All observations recorded as “inaccurate” or “ambiguous” were removed from the climate suitability modelling.

### Home Range Analysis

For species where multivariate accuracy was low (i.e. *Lycorma delicatula*, *Microstegium vimineum*), we performed the standardized Mahalanobis model ‘in reverse’, using North American observations to parameterize the climate envelope, and home range data to test the model outputs. For this variant, we used the final MIC-reduced model where applicable (Table S 4).

### Software

R version 4.3.0 (R Core Team, 2023) was used to build and run the analysis. Species observation data were collected using rgbif (Chamblerlain et al., 2023; Chamberlain and Boettiger, 2017). Spatial calculations and data interoperability were facilitated by terra (Hijmans, 2023), sf (Pebesma, 2018; Pebesma and Bivand, 2023), and data.table (Dowle and Srinivasan, 2023). Plotting was performed using ggplot2 (Wickham, 2016) with tidyterra (Hernangomez, 2023). Internal function benchmarks were calculated using tictoc (Izrailev, 2023). Code was parallelized using the mclapply() function from the parallel package (R Core Team 2023).

## Results

Overall, the standardized Mahalanobis model obtained a high level of accuracy in predicting climate suitability of North American invasions with a relatively high overall multivariate and univariate accuracy (Figure 1; Figure 2; Figures S1-S6). Multivariate accuracy values were below 80% for 6 out of the 23 species, with *Epiphyas postvittana* having 0% accuracy (Table 3).

**Figure 1.**
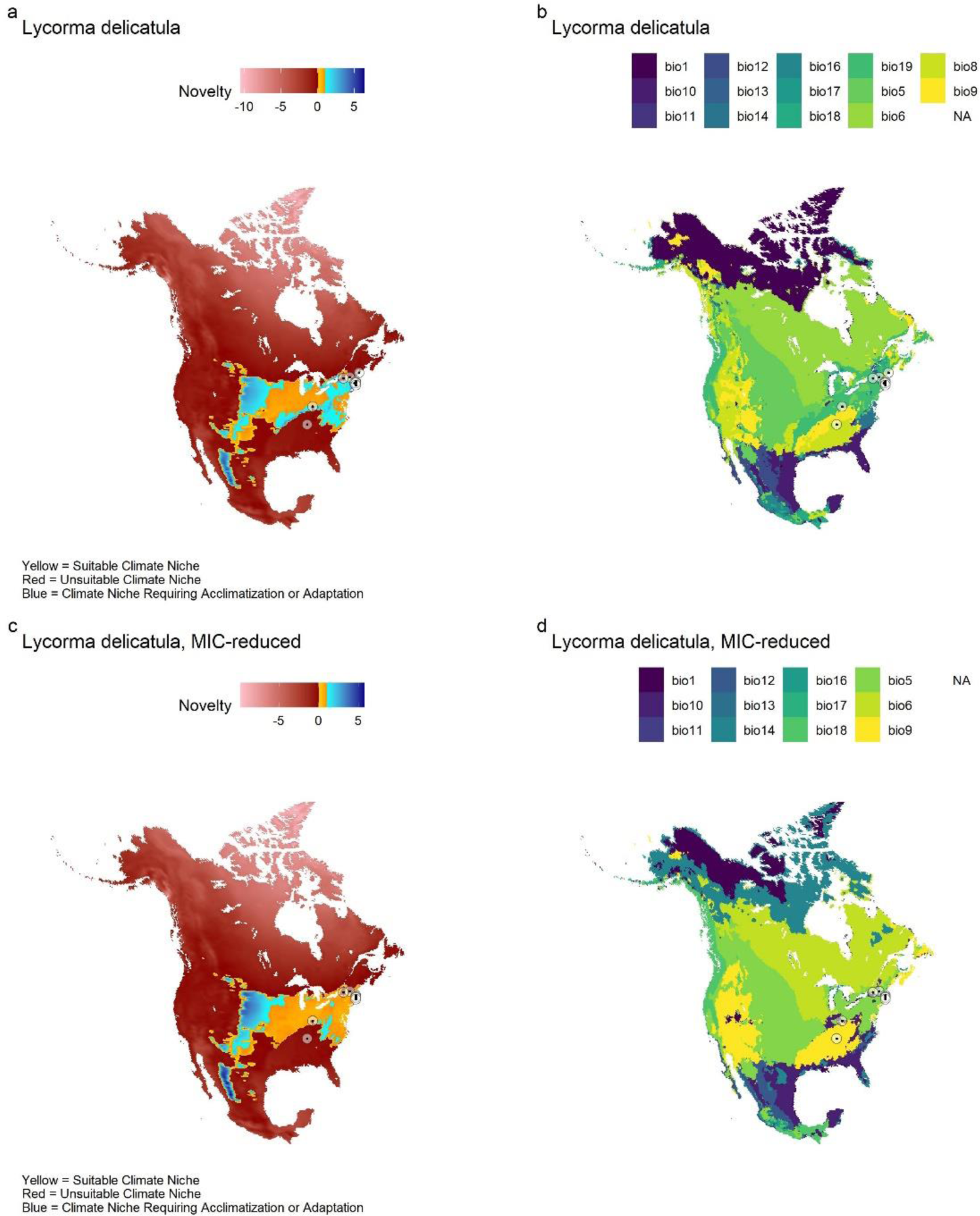
*– Model outputs for Lycorma delicatula* using the full variable set (a, b) and the Most Important Covariate reduced set (MIC-reduced, c, d). a, c) Climate suitability maps showing climate suitability in yellow, univariate climate suitability / type 2 novelty in blue, and unsuitable climate areas in red. b, d) Most important covariates contributing to model results. Black points surrounded by circles indicate observations of *L. delicatula* where the model estimates that the climate is unsuitable. Projection: NAS 1983 Equidistant Conic North America – ESRI:102010.

**Figure 2.**
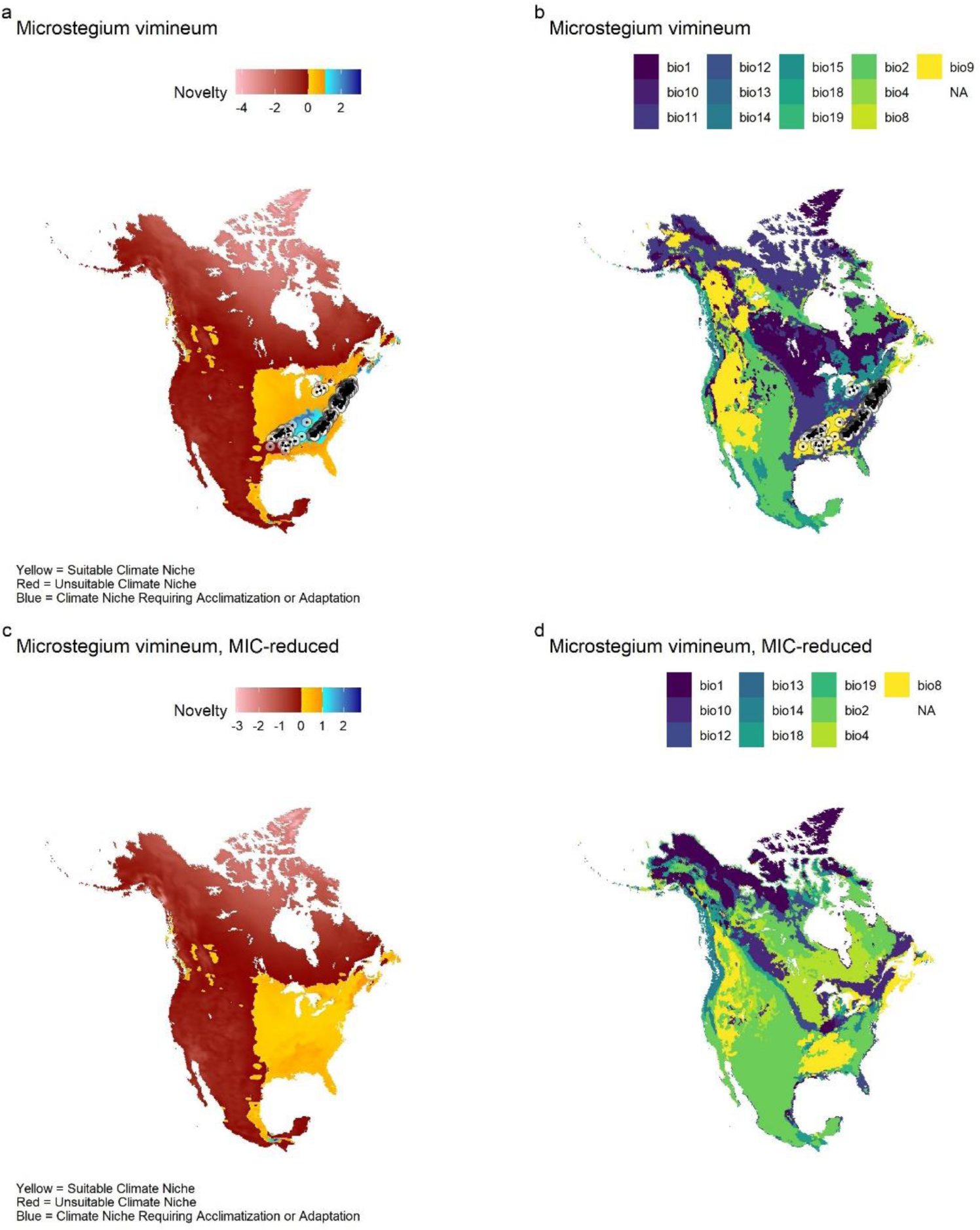
*– Model outputs for Microstegium vimineum* using the full variable set (a, b) and the Most Important Covariate reduced set (MIC-reduced, c, d). a, c) Climate suitability maps showing climate suitability in yellow, univariate climate suitability / type 2 novelty in blue, and unsuitable climate areas in red. b, d) Most important covariates contributing to model results. Black points surrounded by circles indicate observations of *M. vimineum* where the model estimates that the climate is unsuitable. Projection: NAS 1983 Equidistant Conic North America – ESRI:102010.

**Table 3.** – Species distribution model performance for federally regulated plant pest categories in Canada. There were 9 insects and 15 plant species included in the analysis. Overall values are presented as mean ± s.e.m.

| Full Variable Set |  |  |  |  |  |
| --- | --- | --- | --- | --- | --- |
| Category | Species | Multivariate accuracy | Univariate accuracy | Multivariate Error | False Negative |
| Plant | *Aegilops cylindrica* | 95.7% | 95.7% | 4.3% | 4.3% |
| Plant | *Alopecurus myosuroides* | 100% | 100% | 0% | 0% |
| Plant | *Arundo donax* | 100% | 100% | 0% | 0% |
| Plant | *Centaurea iberica* | 83.3% | 83.3% | 16.7% | 16.7% |
| Plant | *Centaurea solstitialis* | 99% | 99% | 1% | 1% |
| Plant | *Crupina vulgaris* | 83.3% | 83.3% | 16.7% | 16.7% |
| Plant | *Echium plantagineum* | 100% | 100% | 0% | 0% |
| Plant | *Eriochloa villosa* | 98.9% | 98.9% | 1.1% | 1.1% |
| Plant | *Microstegium vimineum* | 54.8% | 79.3% | 45.2% | 20.7% |
| Plant | *Paspalum dilatatum* | 97.3% | 97.3% | 2.7% | 2.7% |
| Plant | *Persicaria perfoliata* | 92% | 92.2% | 8% | 7.8% |
| Plant | *Pueraria montana* | 100% | 100% | 0% | 0% |
| Plant | *Senecio inaequidens* | 61.2% | 61.2% | 38.8% | 38.8% |
| Plant | *Zygophyllum fabago* | 78.9% | 78.9% | 21% | 21% |
| Insect | *Cacoecimorpha pronubana* | 100% | 100% | 0% | 0% |
| Insect | *Cydalima perspectalis* | 100% | 100% | 0% | 0% |
| Insect | *Epiphyas postvittana* *** | 0% | 0% | 100% | 100% |
| Insect | *Lobesia botrana* | 57.1% | 57.10% | 42.9% | 42.9% |
| Insect | *Lycorma delicatula* | 60.3% | 97.80% | 39.7% | 2.2% |
| Insect | *Popillia japonica* | 94.6% | 97.00% | 5.4% | 3% |
| Insect | *Rhagoletis cerasi* | 100% | 100% | 0% | 0% |
| Insect | *Sirex noctilio* | 100% | 100% | 0% | 0% |
| Insect | *Tetropium fuscum* | 100% | 100% | 0% | 0% |
| **Plant** | **Overall** | **88.8 ± 4.0%** | **90.6 ± 3.1%** | **11.1 ± 4.0%** | **9.4 ± 3.1%** |
| **Insect** | **Overall** | **79.1 ± 11.5%** | **83.5 ± 11.4%** | **20.9 ± 11.5%** | **16.5 ± 11.4%** |
| **All** | **Overall** | **85.1 ± 5.1%** | **87.9 ± 4.8%** | **14.9 ± 5.1%** | **12.1 ± 4.8%** |
| MIC Reduction |  |  |  |  |  |
| Category | Species | Multivariate accuracy | Univariate accuracy | Multivariate Error | False Negative |
| Plant | *Microstegium vimineum*<sup>a</sup> | 100% | 100% | 0% | 0% |
| Plant | *Senecio inaequidens*<sup>b</sup> | 98.2% | 98.2% | 1.8% | 1.8% |
| Plant | *Zygophyllum fabago*<sup>c</sup> | 84.20% | 84.20% | 15.80% | 15.80% |
| Insect | *Epiphyas postvittana* *** | 0% | 0% | 100% | 100% |
| Insect | *Lobesia botrana*<sup>d</sup> | 85.7% | 100% | 14.3% | 0% |
| Insect | *Lycorma delicatula*<sup>e</sup> | 97.80% | 98.30% | 2.20% | 1.70% |
| **Plant** | **Overall** | **95.1 ± 1.8%** | **95.1 ± 1.8%** | **4.9 ± 1.8%** | **4.9 ± 1.8%** |
| **Insect** | **Overall** | **86.5 ± 10.9%** | **88.4 ± 11.1%** | **13.5 ± 10.9%** | **11.6 ± 11.1%** |
| **All** | **Overall** | **91.7 ± 4.3%** | **92.5 ± 4.4%** | **8.3 ± 4.3%** | **7.5 ± 4.4%** |

\*\*\*For *Epiphys postvittana*, MIC reduction was unable to increase SDM accuracy. Variables removed in MIC reduction were: <sup>a</sup>bio9, bio11, bio15; <sup>b</sup>bio15, bio19; <sup>c</sup>bio2, bio13, bio14, bio18, bio19; <sup>d</sup>bio1, bio8, bio14, bio16, bio18; <sup>e</sup>bio8, bio19.

Using most important covariate (MIC) reduction, it was possible to improve multivariate accuracy to greater than 80% in 5 of the 6 species (Figure 1; Figure 2; Figure S 5; Figure S 6), although it was not possible to improve accuracy for *Epiphyas postvittana*. To illustrate the impact of MIC reduction on climate suitability maps, we looked at *L. delicatula* and *M. vimineum*. For *L. delicatula*, bio19 and bio8 were identified as MICs that led to higher multivariate error. When these variables were eliminated, multivariate accuracy increased from 60.3% to 97.8% (Table 3; Figure 1). For *M. vimineum*, bio9, bio11, and bio15 were identified as MICs that led to higher multivariate error. When these variables were eliminated, multivariate accuracy increased from 54.8% to 100% (Table 3; Figure 2).

Looking at the prediction errors, these tended to cluster near the boundaries of climate suitable regions (Figure 1; Figure S 1a; Figure S 2a,c,g; Figure S 3c,e,g; Figure S 4a; Figure S 5e; Figure S 6a,c,e,g) with some exceptions (Figure 1; Figure S 1a,c; Figure S 2a; Figure S 3c,e; Figure S 4a; Figure S 5a).

Taking all climate suitability estimates together, it becomes possible to identify hotspot areas of North America susceptible to invasion by the CFIA regulated pests analyzed for the present study (Figure 3).

**Figure 3.**
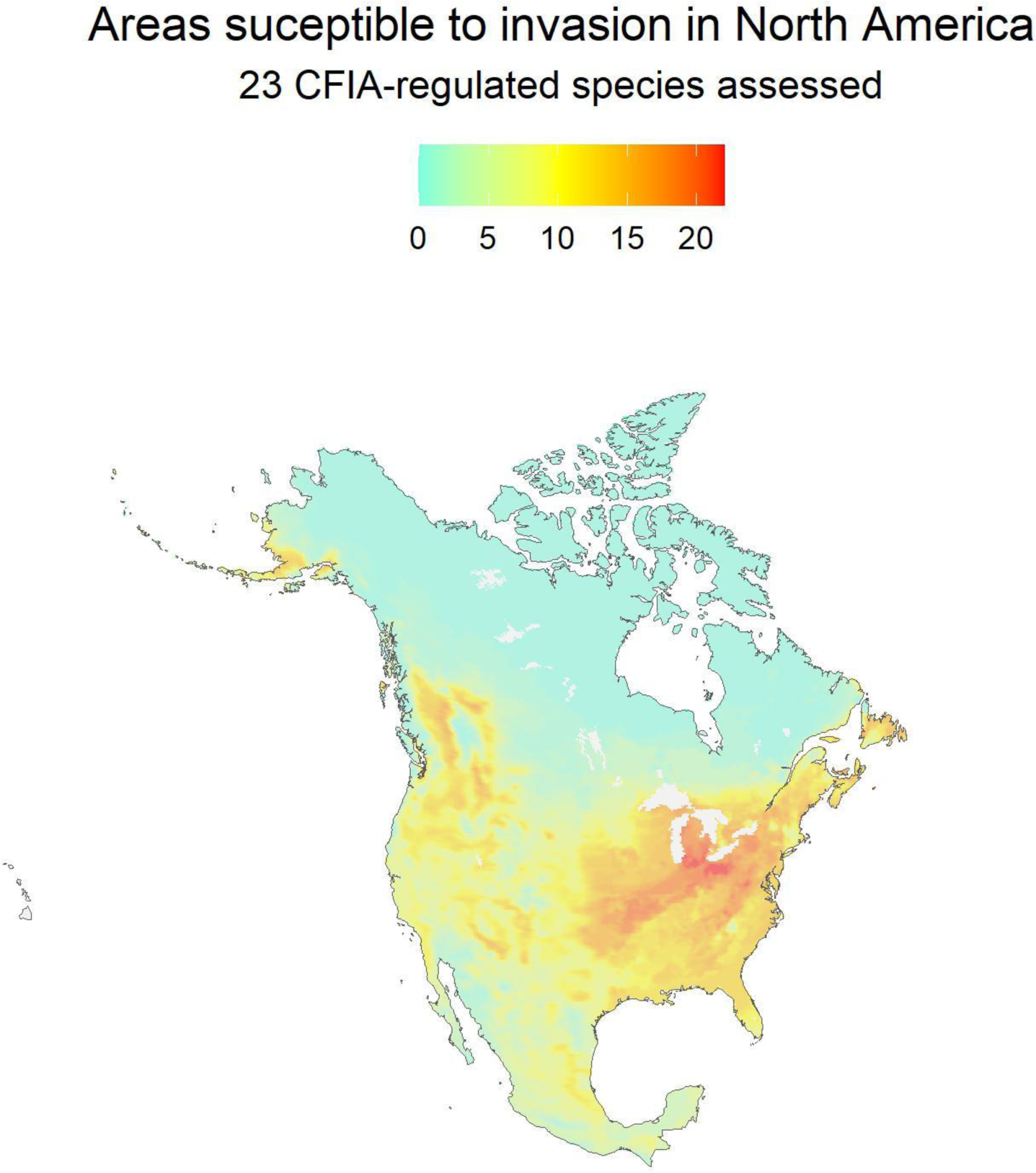
– Areas susceptible to invasion by the 23 CFIA regulated pest species used in the present study. Colour indicates the number of species with climate suitability in a given pixel of the image. Projection: NAS 1983 Equidistant Conic North America – ESRI:102010.

### Home Range Analysis

Accuracy values for the home range analysis were generally quite low, with some exceptions (Table 4). In general, this suggests that the invaded range is generally a smaller subset or partially-shifted subset of the climate envelope of the home range in these species, leading to contradictory results between estimating climate suitability in the invaded range versus the home range.

**Table 4.** – Species distribution model performance when projecting from the invaded North American range into the home range. There were 4 insects and 5 plant species included in the analysis. Overall values are presented as mean ± s.e.m.

| Species | Multivariate accuracy | Univariate accuracy | Multivariate Error | False Negative |
| --- | --- | --- | --- | --- |
| *Aegilops cylindrica* | 90.2% | 91.4% | 9.8% | 8.6% |
| *Alopecurus myosuroides* | 0% | 3.4% | 100% | 96.6% |
| *Arundo donax* | 75% | 76.8% | 25% | 23.2% |
| *Cacoecimorpha pronubana* | 0% | 0% | 100% | 100% |
| *Centaurea iberica* | 0% | 3% | 100% | 97% |
| *Centaurea solstitialis* | 95.9% | 95.9% | 4.1% | 4.1% |
| *Crupina vulgaris* | 0.5% | 19.9% | 99.5% | 80.1% |
| *Cydalima perspectalis* | 94.7% | 94.7% | 5.3% | 5.3% |
| *Echium plantagineum* | 12.3% | 76.9% | 87.7% | 23.1% |
| *Epiphyas postvittana* | 0% | 39.4% | 100% | 60.6% |
| *Eriochloa villosa* | 0% | 0% | 100% | 100% |
| *Lobesia botrana* (MIC reduced) | 0% | 41.7% | 100% | 58.3% |
| *Lycorma delicatula* (MIC reduced) | 0% | 0.8% | 100% | 99.2% |
| *Microstegium vimineum* (MIC reduced) | 49% | 49% | 51% | 51% |
| *Paspalum dilatatum* | 96.1% | 96.1% | 3.9% | 3.9% |
| *Persicaria perfoliata* | 0% | 0% | 100% | 100% |
| *Popillia japonica* | 29.9% | 29.9% | 70.1% | 70.1% |
| *Pueraria montana* | 10.2% | 16.1% | 89.8% | 83.9% |
| *Rhagoletis cerasi* | 0% | 0% | 100% | 100% |
| *Senecio inaequidens* (MIC reduced) | 16.4% | 54.5% | 83.6% | 45.5% |
| *Sirex noctilio* | 0% | 0% | 100% | 100% |
| *Tetropium fuscum* | 0% | 0% | 100% | 100% |
| *Zygophyllum fabago* (MIC reduced) | 12.7% | 46.6% | 87.3% | 53.4% |
| **Overall** | **25.3 ± 7.7%** | **36.4 ± 7.6%** | **74.7 ± 7.7%** | **63.6 ± 7.6%** |

### Sensitivity to iNaturalist data

Due to potential issues that arise in iNaturalist data sources, we looked at the impact on the reference climate range for each species for GBIF data coming from iNaturalist or other sources. In general, we found that the reference climate range was expanded in all species when using iNaturalist occurrence data, though the total number of bioclimatic variables affected differed among species (Table 5).

**Table 5.** – Maximum reference Mahalanobis distance for iNaturalist research-grade observations and non-iNaturalist data, as well as the list of climate variables where the iNaturalist data extended the climate envelope for that variable by more than 20%.

| Species | Maximum Reference Mahalanobis Distance |  | Sensitive Climate Variables |  |
| --- | --- | --- | --- | --- |
|  | Non-iNaturalist Data | iNaturalist Data | Name | Affected / Total |
| *Cacoecimorpha pronubana* | 120 | 178 | bio1, bio5, bio6, bio8, bio9, bio10, bio11, bio12, bio13, bio14, bio16, bio17, bio18, bio19 | 14 / 14 |
| *Cydalima perspectalis* | 356 | 402 | bio13, bio16, bio18 | 3 / 14 |
| *Epiphyas postvittana* | 127 | 142 | bio1, bio5, bio6, bio8, bio9, bio10, bio11, bio12, bio13, bio14, bio16, bio17, bio18, bio19 | 14 / 14 |
| *Lobesia botrana* | 51 | 92 | bio1, bio5, bio6, bio8, bio9, bio10, bio11, bio12, bio13, bio14, bio16, bio17, bio18, bio19 | 14 / 14 |
| *Lycorma delicatula* | 110 | 182 | bio1, bio5, bio6, bio9, bio10, bio11, bio12, bio14, bio17 | 9 / 14 |
| *Popillia japonica* | 32 | 126 | bio1, bio5, bio6, bio8, bio9, bio10, bio11, bio12, bio13, bio14, bio16, bio17, bio18, bio19 | 14 / 14 |
| *Rhagoletis cerasi* | 32 | 110 | bio1, bio5, bio6, bio8, bio9, bio10, bio11, bio12, bio13, bio14, bio16, bio17, bio18, bio19 | 14 / 14 |
| *Sirex noctilio* | 92 | 101 | bio5, bio6, bio9, bio11, bio14, bio17, bio18 | 7 / 14 |
| *Tetropium fuscum* | 70 | 71 | bio10, bio11 | 2 / 14 |
| *Aegilops cylindrica* | 119 | 61 | None | 0 / 13 |
| *Alopecurus myosuroides* | 191 | 223 | bio4 | 1 / 13 |
| *Arundo donax* | 318 | 947 | bio2, bio4, bio12, bio13, bio15, bio19 | 6 / 13 |
| *Centaurea iberica* | 83 | 39 | bio1, bio2, bio8, bio9, bio10, bio12, bio14 | 7 / 13 |
| *Centaurea solstitialis* | 138 | 233 | bio2, bio8, bio9, bio13, bio14, bio15, bio18, bio19 | 8 / 13 |
| *Crupina vulgaris* | 184 | 88 | None | 0 / 13 |
| *Echium plantagineum* | 359 | 572 | bio1, bio2, bio4, bio10, bio12, bio13, bio14, bio15, bio18, bio19 | 10 / 13 |
| *Eriochloa villosa* | 316 | 71 | bio1, bio4, bio8, bio10, bio11, bio12, bio13, bio18 | 8 / 13 |
| *Microstegium vimineum* | 100 | 83 | bio1, bio4, bio8, bio9, bio11, bio12, bio13, bio18 | 8 / 13 |
| *Paspalum dilatatum* | 587 | 432 | bio2, bio4, bio9, bio10, bio12, bio13, bio14, bio18 | 8 / 13 |
| *Persicaria perfoliata* | 82 | 101 | bio1, bio2, bio4, bio9, bio10, bio11, bio12, bio13, bio14, bio15, bio18, bio19 | 12 / 13 |
| *Pueraria montana* | 97 | 127 | bio5, bio6, bio9, bio11 | 4 / 13 |
| *Senecio inaequidens* | 265 | 342 | bio1, bio2, bio4, bio11, bio12, bio13, bio15, bio19 | 8 / 13 |
| *Zygophyllum fabago* | 69 | 113 | bio1, bio8, bio9, bio10, bio11, bio12, bio13, bio18, bio19 | 9 / 13 |

### Spot check of iNaturalist data

iNaturalist observations were inspected / verified for *Alopecurus myosuroides* (336 total) and *Eriochloa villosa* (295 total). For both species, more than 70% of the iNaturalist observations were accurate, however ∼20% of observations were ambiguous and <3% were inaccurate (Table 6).

**Table 6.** – Proportion of iNaturalist observations for two plant species that were accurate identifications, ambiguous identifications, and inaccurate identifications.

| Species | Accurate | Ambiguous | Inaccurate |
| --- | --- | --- | --- |
| *Alopecurus myosuroides* | 77.5% | 20.5% | 1.8% |
| *Eriochloa villosa* | 74.9% | 22.4% | 2.7% |

## Discussion

The standardized Mahalanobis distance as presented by Mesgaran et al. (2014) achieved a high level of accuracy for predicting the climate suitability of invasive species in North America (Figure 1; Figure 2; Figures S1-S6; Table 3), supporting our objective to demonstrate the utility of the model for predicting potential invasion sites. Variable reduction through MIC analysis was able to achieve some improvements to model accuracy. Overall, this suggests that the standardized Mahalanobis distance model, despite harnessing the covariance of climate variables, is still sensitive to being over-constrained with too many variables.

The model itself captures variability metrics for climate-suitable areas, with positive values closer to 0 indicating areas climatically similar to the centroid of the climate envelope, and values closer to 1 indicating potential constraints on the climate envelope including physical barriers, competition, or marginal climates. Meanwhile, the type 2 novelty projections provide a quantitative measure of how climatically different an area is from the reference climate within the multivariate climate envelope, which may be useful in prioritizing surveillance in areas outside of strictly climate-suitable areas.

When looking at prediction errors, these tended to be clustered near the edge of climate-suitable areas (Figure 1; Figure 2; Figures S1-S6). These ‘edge’ errors could be due to a number of factors, including: spatial aggregation of climate data and matched species presence data, underestimation of the climate envelope, over-constraining the model, or an actual shift in climate envelope. The MIC reduction analysis supports the idea that over-constraining the model was responsible in most cases (Figure 1; Figure 2; Figure S 5; Figure S 6). However, *Epiphyas postvittana* was an exception with respect to prediction error, where MIC analysis was unable to improve the model’s ability to accurately capture the North American invasion points. Hill et al. (2017) showed that the climatic niche of *E. postvittana* expanded by over 10% of its original size during invasion. Since we only used the non-North American range for assessing climate suitability of *E. postvittana*, niche expansion of the North American invasive is a possible explanation for why the standardized Mahalanobis distance was unable to accurately identify areas of its North American invasion. Adaptation within the invaded range may also explain the inability of the model to identify areas of the North American invasion as being climate suitable, as Morey et al. (2013) demonstrate the potential impact of selection on expanding the range limits of *E. postvittana*.

The model performed quite poorly when reversing the direction and applying the model from the invaded North American range to the home range of the species (Table 4). The primary reason for the poor performance when reversing the model is due to the North American invasions occupying a subset of the reference climate envelope, such that the reference observations lie outside or within only a portion of the climate envelope calculated from the North American observation points. We expect these results (Broennimann et al., 2007; Liu et al., 2020), especially for ongoing invasions, for reasons including time to spread throughout a pest’s potential invasion range, part of the home range climate envelope not existing within the invaded range, and potential biotic constraints preventing a pest from spreading to its full climate envelope within the invaded range (Walder et al., 2019). Additional explanations include potential founder effects narrowing the climate envelope of the invading population (Liu et al., 2020), niche shifts (Broennimann et al., 2007; Bujan et al., 2021; van Boheeman et al., 2019) or adaptation (e.g. Bujan et al., 2021; van Boheeman et al., 2019).

### Limitations and Assumptions

The predictions of climate suitability for the standardized Mahalanobis SDM are limited by the observed climate envelope. The limitations and assumptions of this method include:

- **Data Quality for Training and Testing**: Many of the data points used for training and testing the model involved research grade observations from iNaturalist that were contributed to GBIF. Some of these observations may be misidentifications (Table 6), since a research grade rating is granted to any iNaturalist observation with at least 2 confirmations of the species ID. As such, this introduces a level of potential error into both the projected climate suitability as well as the calculation of model accuracy due to the influence of iNaturalist observations on climate envelope calculations (Table 5). Additionally, there are some instances where the photos in iNaturalist data were not taken at the location indicated (Mooij, pers. comm.), which could introduce errors in the both the model testing presented here, and in constructing the multivariate climate envelope with the model.
- **Sample selection:** Occurrence records are often recorded in disturbed areas, areas with higher human density, and sites that are easier to sample, which affects citizen science, herbarium, and laboratory diagnostic data.
- **Observed Climate Envelope:** If too little of the climate envelope is captured in the presence data, the predictions will underestimate the full climatic breadth suitable to the species.
- **Interpretation of Type 1 Novel Environments:** This technique assumes that Type 1 Novel environments are not climate-suitable – such environments may be climate-suitable if the observed envelope is smaller than the actual climate envelope or if the species is capable of adapting to the Type 1 Novel environment. Note that niche shifts during invasion have been observed (Broennimann et al., 2007).
- **Interpretation of Type 2 Novel Environments:** Type 2 Novel environments may or may not be climate-suitable depending on whether the Type 2 Novel environment falls within the actual climate envelope or whether the species can acclimatize or adapt to the novel combinations of climatic variables. This makes it difficult to assess if a species observation in a Type 2 Novel environment is actually novel, or the result of acclimatization or adaptation.
- **Number of Climatic Variables:** More climate variables used in the SDM will reduce the size of the observed climate envelope. Since the SDM is correlative, this could lead to an artificial limitation on the predicted area of climatic suitability (Table 3; Figure 1; Figure 2; Figure S 5; Figure S 6), especially if any of the climate variables are not biologically relevant. Alternatively, selecting too few climate variables with this method may overestimate the potential climate suitable range for a species.

### Conclusions and Recommendations

The standardized Mahalanobis distance calculation, developed by Mesgaran et al. (2014) as a diagnostic of SDMs, but implemented here as an SDM, is an effective means to identify areas of potential invasion of North America by invasive plant pests. The model can be run quickly (< 20 min) on a PC equipped with 16 GB RAM and 4-core processor, with minimal biological knowledge of the organism, making the SDM an effective tool for rapidly determining the potential area of invasion for guiding early detection and rapid response activities until additional biological information is available.

Based on our analysis, we recommend that the SDM be used with a tailored, but complete set of climate variables. The initial climate variable sets used in the present study represent a ‘first pass’ without tailoring the analysis, while the MIC reduction analysis shows the importance of paring down the climate variables to improve projections. In cases where data are more limited, such as can be in formal pest risk assessments, interpreting the results may need to account for an underestimation of the climate envelope, and could be augmented by adding uncertainties in the reference climate envelope based on empirical data. For example, augmenting the lower limit for bio6 (mean daily minimum air temperature of the coldest month) by 1 to 2 °C if the species is known to be more cold tolerant than the observational data suggest could reduce the impact of low numbers of observations.

We also note that this model is not limited to climatic variables. For example, in the case of plants, soil types could influence the potential distribution of an invasive species, and this model could incorporate soil information to generate estimates of the potential species distribution. However, such a use case would require additional testing.

## Acknowledgments

We would like to thank Karen Castro and Alexandre Blain for their input on climate variables to use for plant species, and Graham Thurston for confirming whether *Cydalima perspectalis* observations from Canadian Food Inspection Agency were from nurseries or outdoor observations. We would like to thank the Canadian Food Inspection Agency laboratories, specifically the Saskatoon Laboratory – Seed Science and Technology, the Ottawa Plant Laboratory – Genotyping and Botany, and the Ottawa Plant Laboratory – Entomology for contributing diagnostic data to test the species distribution model. We would like to thank Graham Thurston, Marie-Claude Gagnon, and Diana Mooij for providing reviews of the manuscript. We would also like to thank Wendy Asbil, Erin LeClair, Arvind Vasudevan, Javier Maldonado, and Sarah Brearey for providing reviews of the CFIA data.

## Contributions

J Stinziano conceived of the study and ran the analysis. A Charron reviewed the research grade iNaturalist reports to confirm species identifications. J Stinziano and M Damus designed the study. J Stinziano wrote the manuscript with input from A Charron and M Damus.

## Supplementary Information

### Supplementary Tables

**Table S 1.** – Value ranges of bioclimatic variables used to calculate the reference climate for the species distribution model. Data shown are for insects.

|  | *Cacoecimorpha pronubana* |  | *Cydalima perspectalis* |  | *Epiphyas postvittana* |  | *Lobesia botrana* |  | *Lycorma delicatula* |  | *Popillia japonica* |  | *Rhagoletis cerasi* |  | *Sirex noctilio* |  | *Tetropium fuscum* |  |
| --- | --- | --- | --- | --- | --- | --- | --- | --- | --- | --- | --- | --- | --- | --- | --- | --- | --- | --- |
|  | min | max | min | max | min | max | min | max | min | max | min | max | min | max | min | max | min | max |
| bio1 | 3.69 | 18.26 | -0.21 | 26.23 | 8.16 | 20.86 | 5.84 | 18.62 | 7.14 | 18.75 | 0.29 | 17.50 | -0.54 | 14.15 | -0.69 | 18.20 | -0.88 | 12.37 |
| bio5 | 14.92 | 30.99 | 13.32 | 33.75 | 14.08 | 28.09 | 19.60 | 32.92 | 18.65 | 32.90 | 15.07 | 30.20 | 14.52 | 29.92 | 12.36 | 31.66 | 13.43 | 25.48 |
| bio6 | -12.29 | 11.21 | -19.61 | 19.15 | 0.01 | 13.68 | -7.94 | 13.08 | -17.21 | 3.97 | -14.51 | 13.84 | -19.50 | 2.47 | -14.75 | 8.01 | -20.41 | 3.11 |
| bio8 | 0.90 | 19.02 | -5.18 | 28.01 | 5.24 | 23.80 | 2.32 | 20.76 | 5.98 | 27.55 | 3.68 | 24.93 | -4.58 | 20.67 | -2.95 | 20.05 | -7.83 | 18.79 |
| bio9 | -4.38 | 25.03 | -10.49 | 27.17 | 5.24 | 19.63 | -1.64 | 25.28 | -7.95 | 13.87 | -9.24 | 20.40 | -12.80 | 21.84 | -5.78 | 25.43 | -12.07 | 19.95 |
| bio10 | 12.80 | 25.06 | 8.76 | 28.09 | 12.83 | 24.42 | 14.64 | 25.37 | 15.57 | 28.44 | 9.82 | 26.38 | 8.92 | 24.12 | 8.85 | 25.45 | 8.53 | 19.95 |
| bio11 | -5.96 | 13.54 | -10.49 | 24.43 | 3.21 | 16.76 | -3.18 | 14.22 | -7.95 | 9.98 | -9.24 | 14.71 | -15.08 | 7.02 | -11.30 | 11.83 | -15.75 | 5.97 |
| bio12 | 478 | 2261 | 378 | 4505 | 595 | 2169 | 513 | 1905 | 267 | 3084 | 796 | 2546 | 423 | 1909 | 290 | 3184 | 488 | 2901 |
| bio13 | 58.35 | 377 | 47.59 | 836 | 60.75 | 366 | 62.24 | 343 | 55.77 | 495 | 85.40 | 531 | 45.41 | 241 | 37.88 | 397 | 55.90 | 347 |
| bio14 | 0.94 | 100 | 0.21 | 217 | 17.37 | 107 | 1.32 | 111 | 1.30 | 185 | 26.81 | 133 | 4.63 | 110 | 3.50 | 136 | 15.79 | 160 |
| bio16 | 160 | 951 | 122 | 2074 | 168 | 906 | 178 | 825 | 150 | 1103 | 250 | 1205 | 124 | 695 | 108 | 1101 | 161 | 925 |
| bio17 | 13.11 | 317 | 2.78 | 758 | 93.74 | 334 | 11.84 | 381 | 6.02 | 610 | 98.37 | 456 | 37.88 | 367 | 27.61 | 4401 | 57.47 | 514 |
| bio18 | 28.28 | 578 | 8.29 | 2028 | 124 | 547 | 24.11 | 670 | 150 | 1068 | 154 | 1205 | 40.26 | 695 | 32.42 | 587 | 143 | 747 |
| bio19 | 114 | 841 | 9.88 | 1028 | 137 | 705 | 93.37 | 768 | 6.32 | 850 | 137 | 587 | 37.88 | 466 | 69.97 | 1029 | 67.88 | 925 |

**Table S 2.** – Value ranges of bioclimatic variables used to calculate the reference climate for the species distribution model. Data shown are for plants.

|  | *Aegilops*<br>*cylindrica* |  | *Alopecurus*<br>*myosuroides* |  | *Arundo donax* |  | *Centaurea*<br>*iberica* |  | *Centaurea*<br>*solstitialis* |  | *Crupina*<br>*vulgaris* |  | *Echium*<br>*plantagineum* |  |
| --- | --- | --- | --- | --- | --- | --- | --- | --- | --- | --- | --- | --- | --- | --- |
|  | min | max | min | max | min | max | min | max | min | max | min | max | min | max |
| bio1 | -2.40 | 28.59 | -0.71 | 22.43 | -1.98 | 28.89 | 1.63 | 23.66 | -2.40 | 28.27 | -2.40 | 18.89 | -1.55 | 22.54 |
| bio2 | 2.49 | 14.83 | 1.30 | 14.78 | 0.92 | 15.81 | 4.19 | 12.29 | 1.57 | 13.88 | 1.56 | 12.26 | 0.92 | 13.66 |
| bio4 | 125 | 1147 | 203 | 1098 | 22.68 | 944 | 430 | 1245 | 245 | 1052 | 399 | 1025 | 91.93 | 892 |
| bio8 | -5.47 | 28.58 | -2.88 | 28.98 | -7.27 | 31.78 | -4.32 | 20.12 | -4.36 | 29.14 | -5.18 | 25.29 | -5.81 | 29.01 |
| bio9 | -11.90 | 34.03 | -8.89 | 31.93 | -6.89 | 36.12 | -8.25 | 30.30 | -10.44 | 31.10 | -10.44 | 27.75 | -8.52 | 29.64 |
| bio10 | 6.99 | 34.03 | 8.31 | 34.11 | 7.77 | 36.12 | 11.50 | 30.33 | 6.99 | 33.71 | 6.99 | 27.75 | 7.69 | 29.64 |
| bio11 | -11.90 | 26.91 | -11.03 | 16.35 | -12.81 | 26.31 | -9.85 | 16.24 | -11.81 | 21.84 | -11.81 | 13.25 | -11.26 | 18.09 |
| bio12 | 64.61 | 2409 | 35.12 | 2294 | 1.82 | 4841 | 120 | 2299 | 89.25 | 3013 | 234 | 1972 | 23.38 | 2430 |
| bio13 | 15.97 | 418 | 7.28 | 302 | 0.43 | 853 | 20.66 | 302 | 18.55 | 431 | 26.48 | 343 | 5.10 | 382 |
| bio14 | 0.01 | 141 | 0.05 | 148 | 0.00 | 293 | 0.20 | 134 | 0.20 | 133 | 0.51 | 124 | 0.00 | 151 |
| bio15 | 11.38 | 101 | 7.55 | 121 | 9.46 | 160 | 16.92 | 113 | 8.95 | 113 | 10.28 | 87.77 | 9.26 | 115 |
| bio18 | 0.15 | 984 | 0.23 | 746 | 0.00 | 2175 | 0.70 | 754 | 1.33 | 746 | 8.04 | 550 | 0.31 | 761 |
| bio19 | 24.84 | 719 | 19.46 | 736 | 0.56 | 987 | 45.03 | 792 | 33.76 | 1013 | 46.25 | 768 | 10.93 | 886 |

**Table S 3.** – Value ranges of bioclimatic variables used to calculate the reference climate for the species distribution model. Data shown are for plants.

|  | *Eriochloa villosa* |  | *Microstegium vimineum* |  | *Paspalum dilatatum* |  | *Persicaria perfoliata* |  | *Pueraria montana* |  | *Senecio inaequidens* |  | *Zygophyllum fabago* |  |
| --- | --- | --- | --- | --- | --- | --- | --- | --- | --- | --- | --- | --- | --- | --- |
|  | min | max | min | max | min | max | min | max | min | max | min | max | min | max |
| bio1 | -0.18 | 22.88 | 2.91 | 25.13 | 3.76 | 27.60 | -0.15 | 24.19 | -10.36 | 27.53 | -4.37 | 18.49 | 1.81 | 19.35 |
| bio2 | 2.23 | 11.23 | 1.88 | 11.36 | 1.00 | 13.77 | 2.45 | 10.39 | 1.31 | 13.78 | 1.62 | 14.62 | 2.25 | 14.24 |
| bio4 | 408 | 1636 | 177 | 1267 | 23.49 | 846 | 264 | 1521 | 39.28 | 1129 | 222 | 839 | 362 | 1343 |
| bio8 | 9.34 | 27.24 | -0.95 | 28.13 | -1.07 | 31.28 | 2.57 | 29.02 | 0.09 | 28.81 | -5.47 | 23.83 | -1.72 | 22.56 |
| bio9 | -22.55 | 20.99 | -11.62 | 23.34 | -2.20 | 27.47 | -18.88 | 23.34 | -19.12 | 29.59 | -13.71 | 24.97 | -5.69 | 29.35 |
| bio10 | 15.59 | 27.86 | 12.80 | 28.13 | 12.38 | 32.14 | 15.80 | 29.03 | 0.09 | 31.69 | 5.29 | 25.16 | 11.33 | 30.42 |
| bio11 | -22.55 | 17.01 | -11.62 | 22.53 | -6.08 | 25.23 | -20.90 | 19.58 | -19.74 | 26.36 | -13.98 | 13.44 | -11.71 | 13.80 |
| bio12 | 518 | 2542 | 527 | 4732 | 103 | 4737 | 581 | 4547 | 394 | 4842 | 292 | 3825 | 140 | 1280 |
| bio13 | 61.37 | 475 | 80.82 | 813 | 18.93 | 815 | 80.47 | 701 | 68.40 | 897 | 39.76 | 472 | 19.69 | 152 |
| bio14 | 5.38 | 130 | 3.68 | 242 | 0.66 | 216 | 9.36 | 216 | 0.61 | 276 | 1.26 | 180 | 0.00 | 64.07 |
| bio15 | 18.43 | 108 | 12.65 | 126.14 | 9.52 | 130 | 10.20 | 102 | 10.41 | 141 | 7.55 | 73.11 | 11.42 | 103 |
| bio18 | 154 | 1102 | 227 | 2250 | 3.36 | 2071 | 226 | 1952 | 127 | 2250 | 20.37 | 875 | 0.73 | 326 |
| bio19 | 24.82 | 715 | 14.70 | 1023 | 4.37 | 852 | 31.28 | 915 | 4.29 | 1161 | 23.11 | 1140 | 35.86 | 368 |

**Table S 4.**
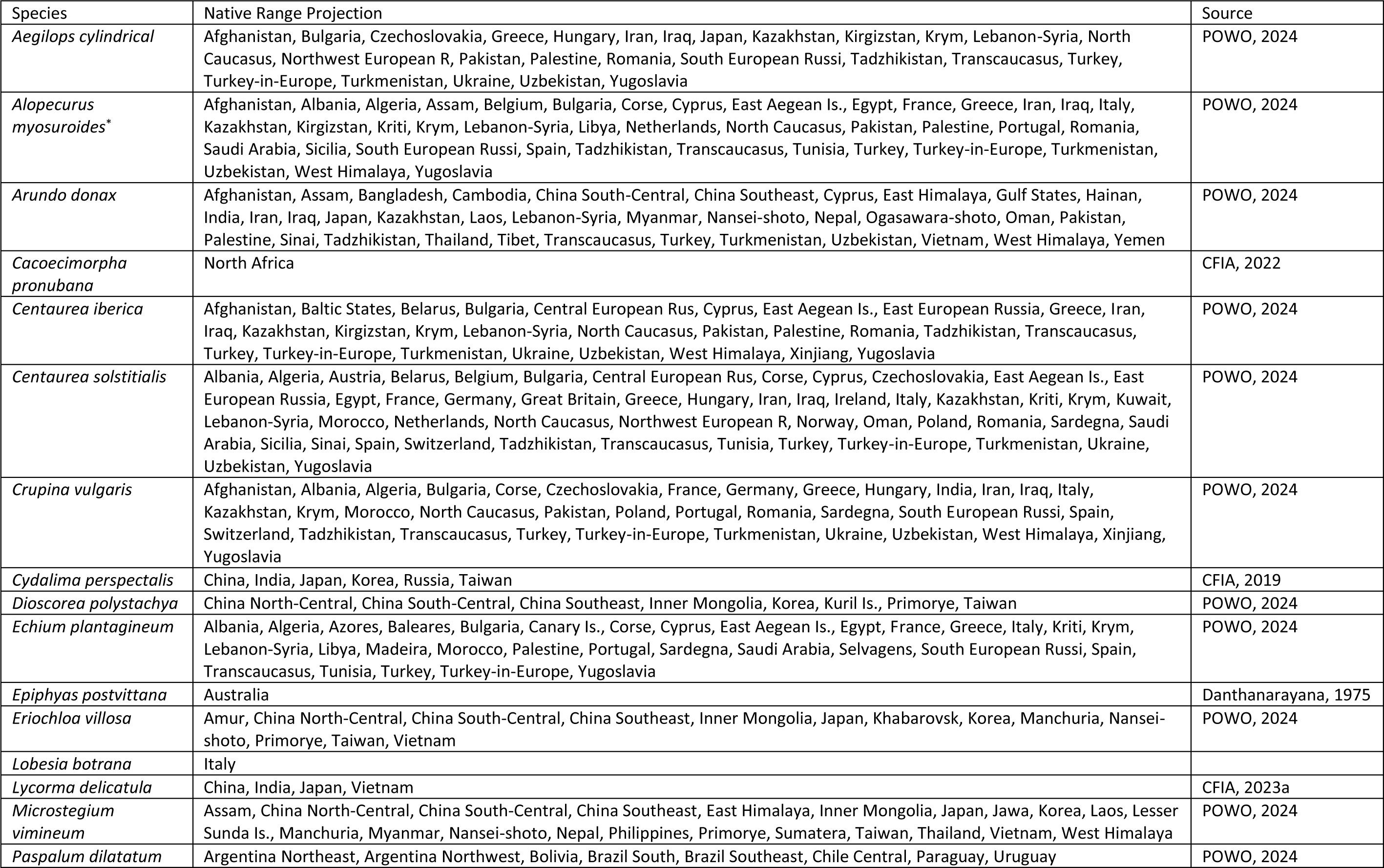
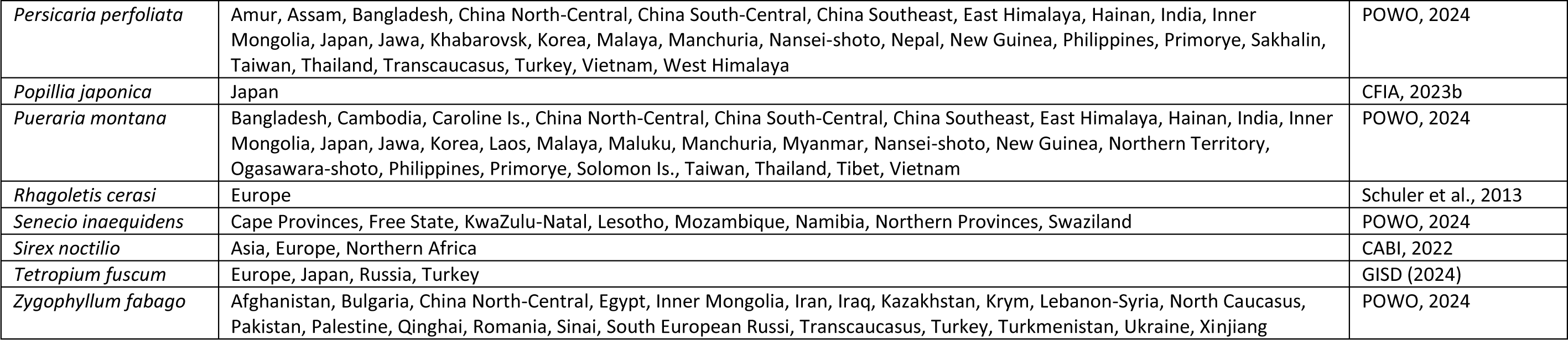
– Native ranges used for the home range analysis.

### Supplementary Figures

**Figure S 1.**
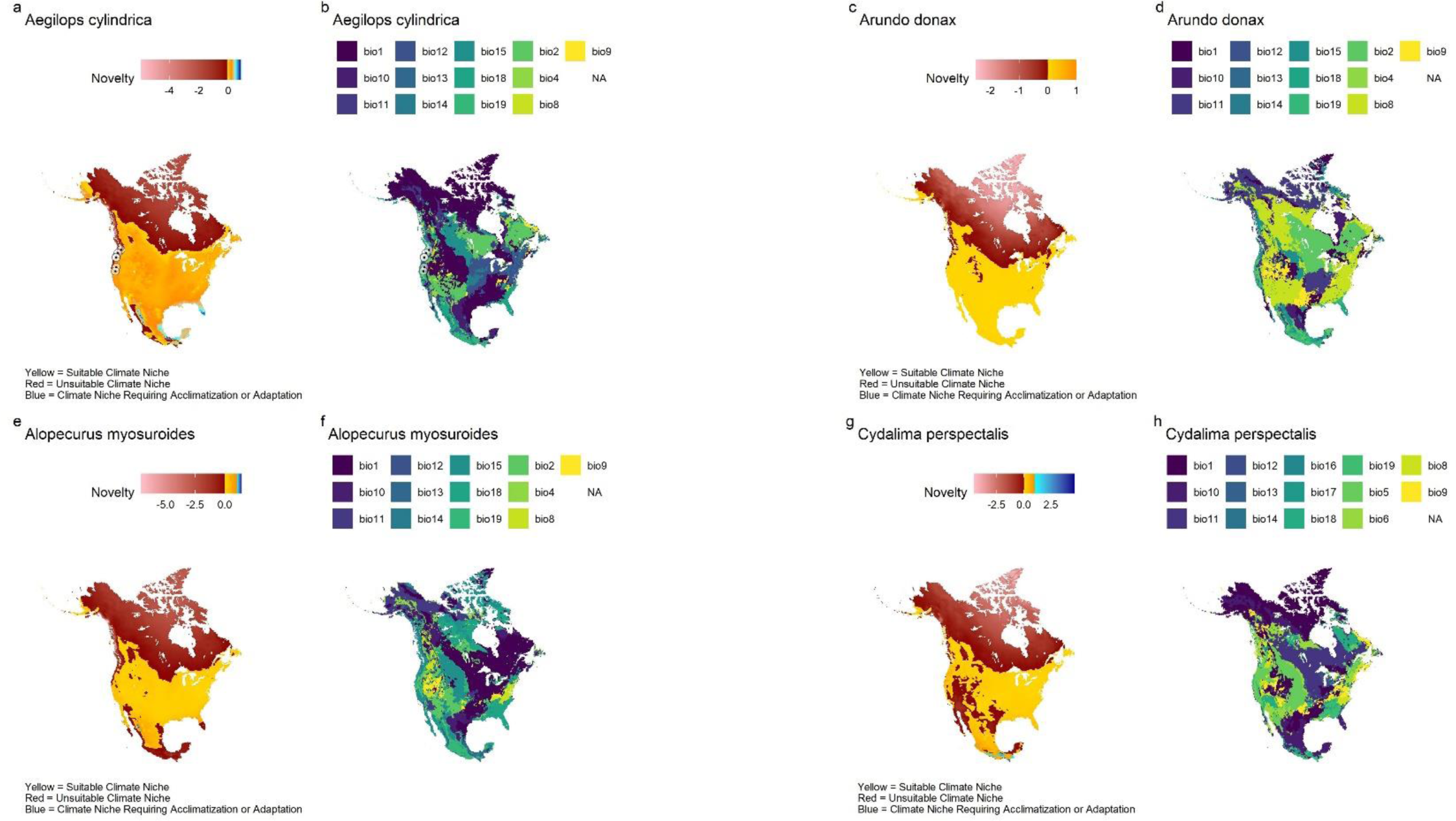
*– Model outputs for Aegilops cylindrica* (a,b), *Arundo donax* (c,d), *Alopecurus myosuroides* (e,f), and *Cydalima perspectalis* (g,h). a, c, e, g) Climate suitability maps showing climate suitability in yellow, univariate climate suitability / type 2 novelty in blue, and unsuitable climate areas in red. b, d, f, h) Most important covariates contributing to model results. Black points surrounded by circles indicate observations of each species where the model estimates that the climate is unsuitable. Projection: NAS 1983 Equidistant Conic North America – ESRI:102010.

**Figure S 2.**
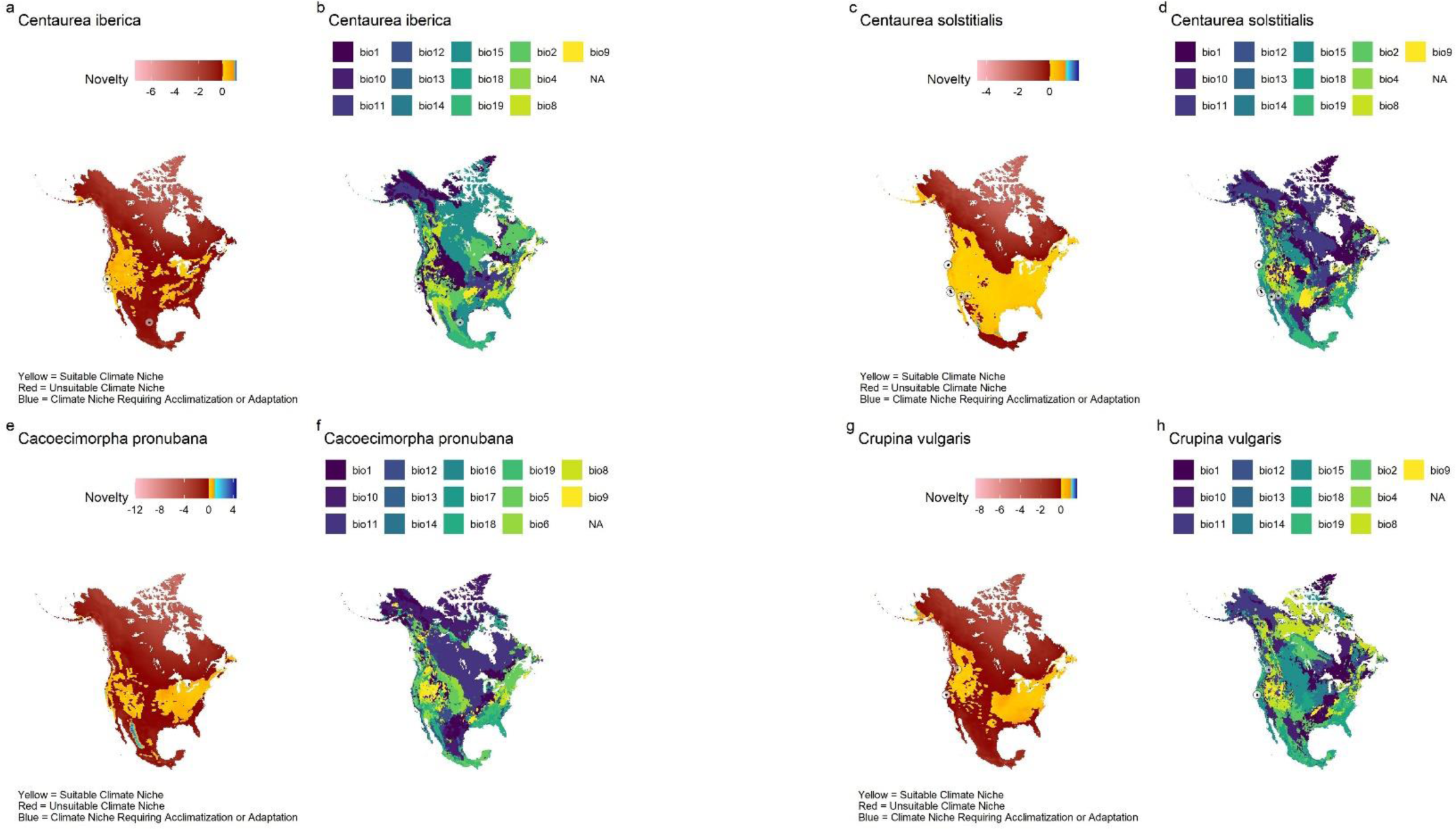
*– Model outputs for Centaurea iberica* (a,b), *Centaurea solstitialis* (c,d), *Cacoecimorpha pronubana* (e,f), and *Crupina vulgaris* (g,h). a, c, e, g) Climate suitability maps showing climate suitability in yellow, univariate climate suitability / type 2 novelty in blue, and unsuitable climate areas in red. b, d, f, h) Most important covariates contributing to model results. Black points surrounded by circles indicate observations of each species where the model estimates that the climate is unsuitable. Projection: NAS 1983 Equidistant Conic North America – ESRI:102010.

**Figure S 3.**
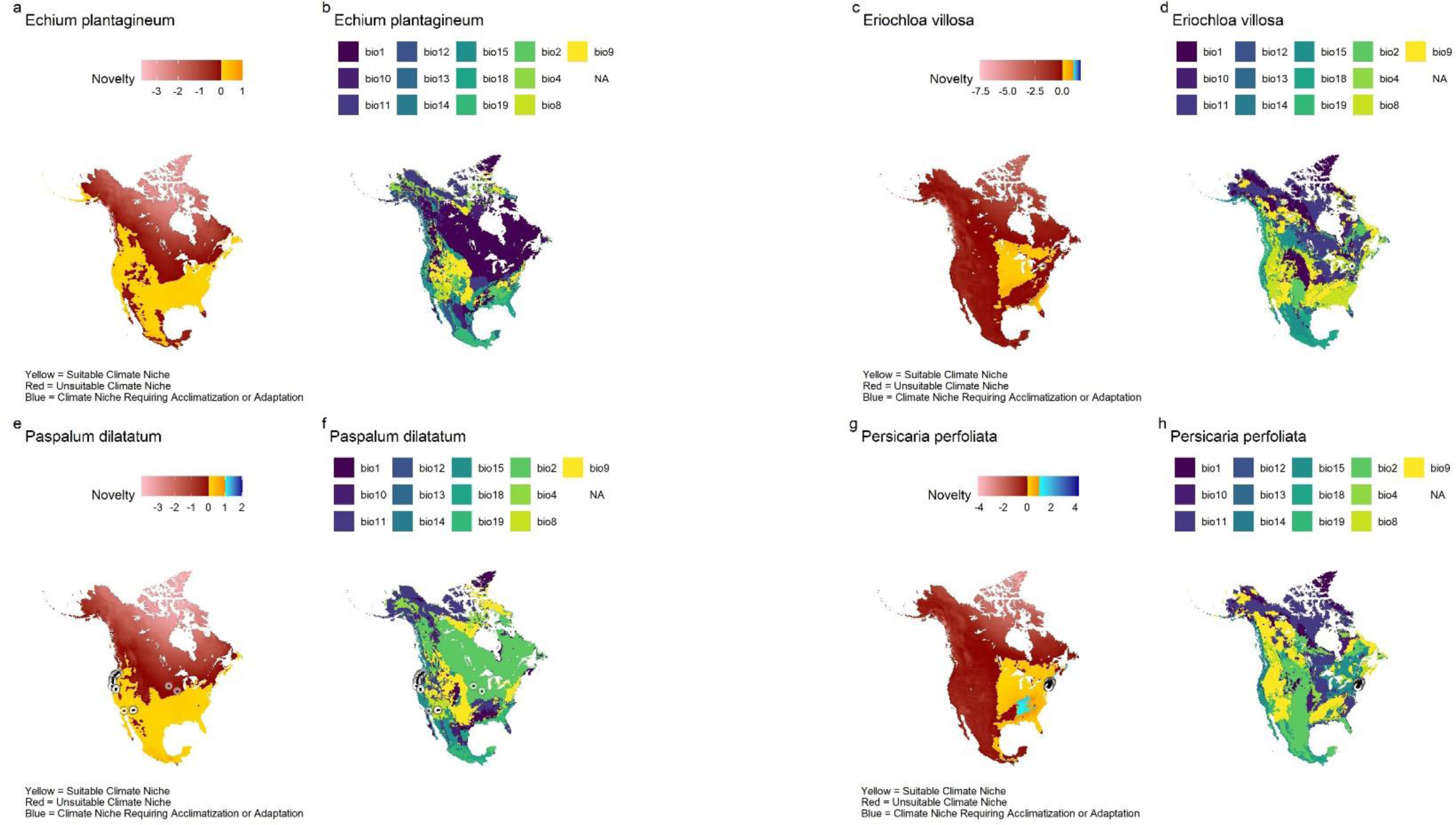
*– Model outputs for Echium plantagineum* (a,b), *Eriochloa villosa* (c,d), *Paspalum dilatatum* (e,f), and *Persicaria perfoliata* (g,h). a, c, e, g) Climate suitability maps showing climate suitability in yellow, univariate climate suitability / type 2 novelty in blue, and unsuitable climate areas in red. b, d, f, h) Most important covariates contributing to model results. Black points surrounded by circles indicate observations of each species where the model estimates that the climate is unsuitable. Projection: NAS 1983 Equidistant Conic North America – ESRI:102010.

**Figure S 4.**
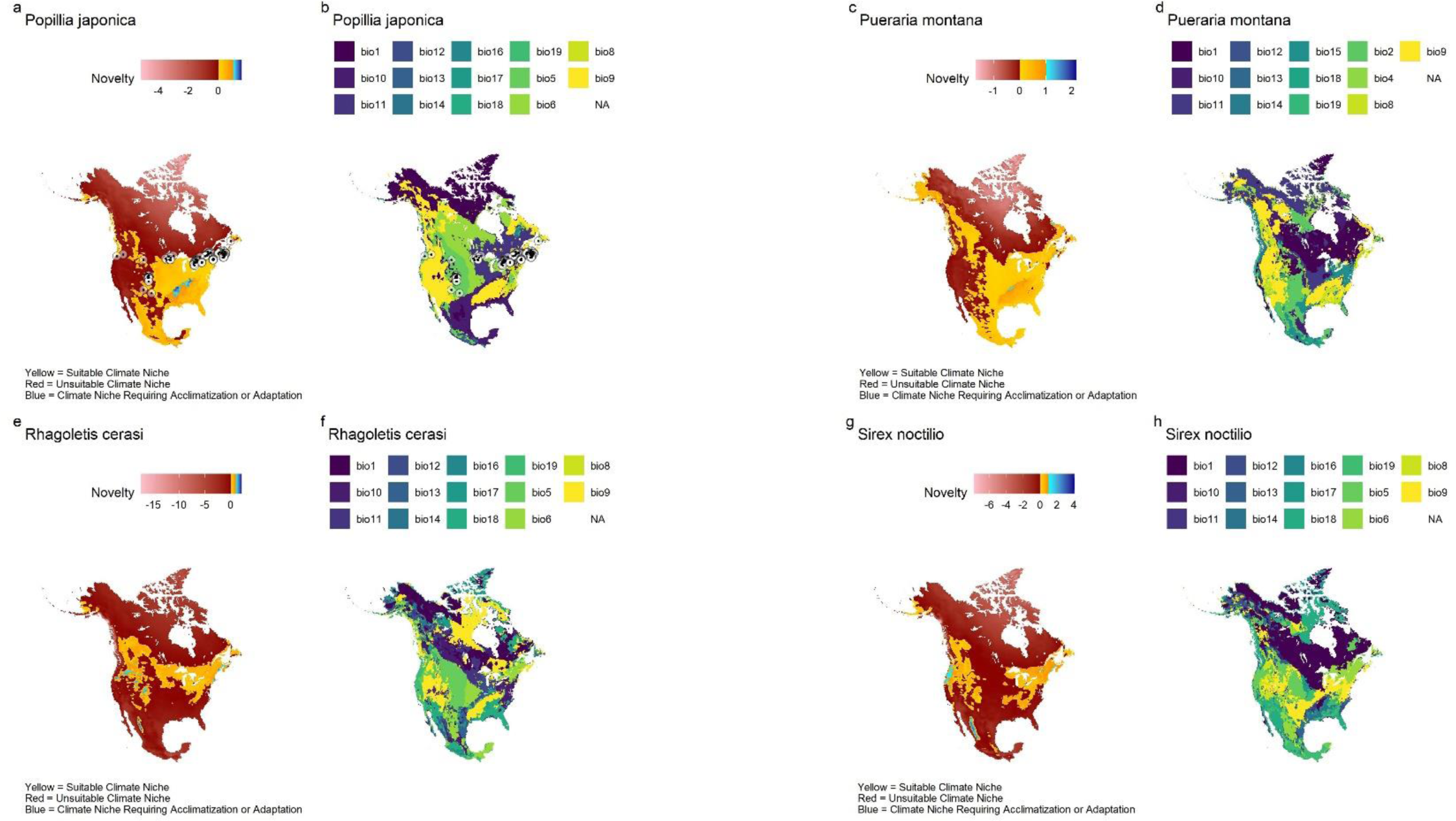
*– Model outputs for Popillia japonica* (a,b), *Pueraria montana* (c,d), *Rhagoletis cerasi* (e,f), and *Sirex noctilio* (g,h). a, c, e, g) Climate suitability maps showing climate suitability in yellow, univariate climate suitability / type 2 novelty in blue, and unsuitable climate areas in red. b, d, f, h) Most important covariates contributing to model results. Black points surrounded by circles indicate observations of each species where the model estimates that the climate is unsuitable. Projection: NAS 1983 Equidistant Conic North America – ESRI:102010.

**Figure S 5.**
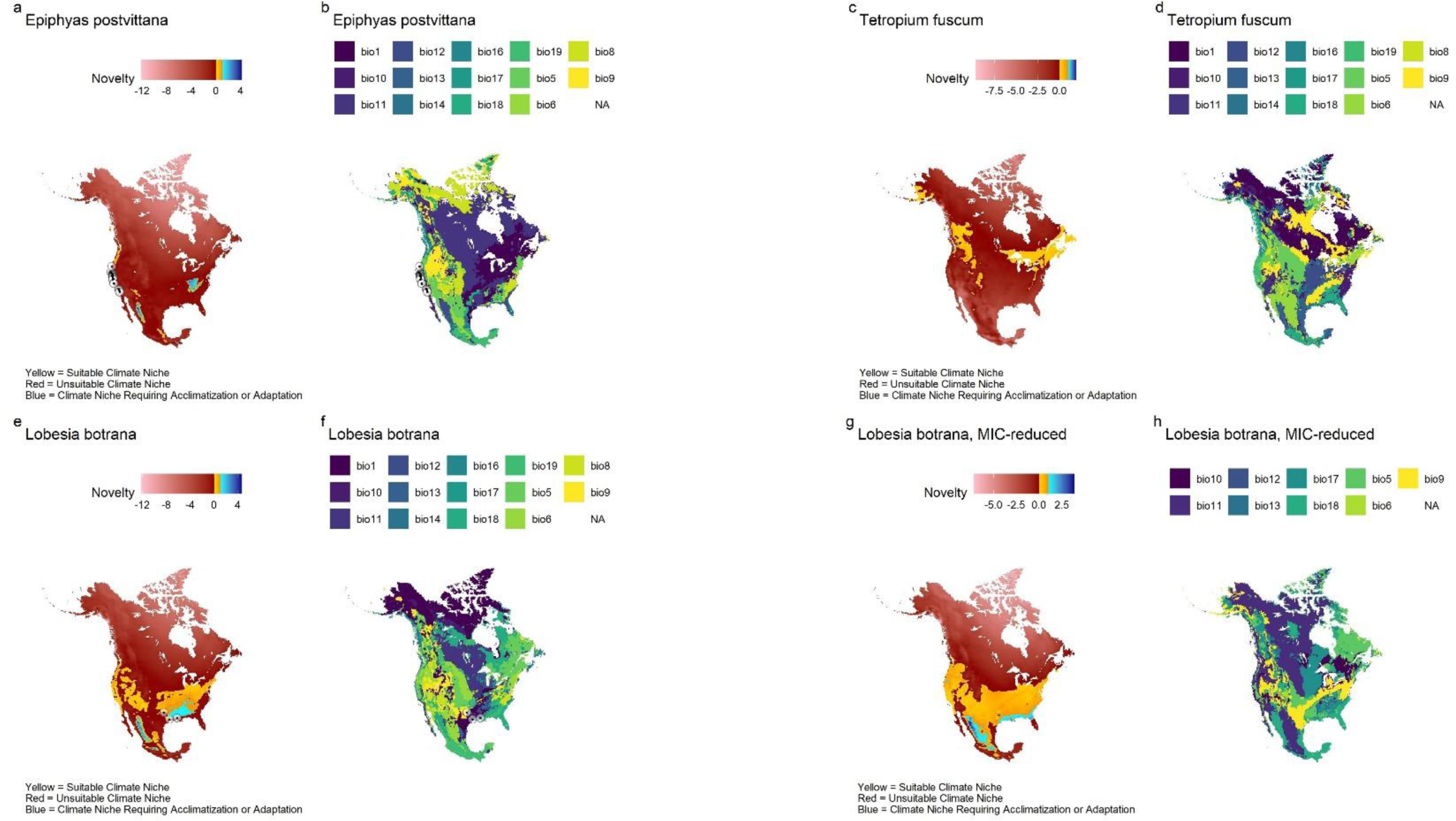
– *Model outputs for Epiphyas postvittana* (a,b), *Tetropium fuscum* (c,d), and *Lobesia botrana* with the full variable set (e,f) and with the MIC-reduced output (g,h). a, c, e, g) Climate suitability maps showing climate suitability in yellow, univariate climate suitability / type 2 novelty in blue, and unsuitable climate areas in red. b, d, f, h) Most important covariates contributing to model results. Black points surrounded by circles indicate observations of each species where the model estimates that the climate is unsuitable. Projection: NAS 1983 Equidistant Conic North America – ESRI:102010.

**Figure S 6.**
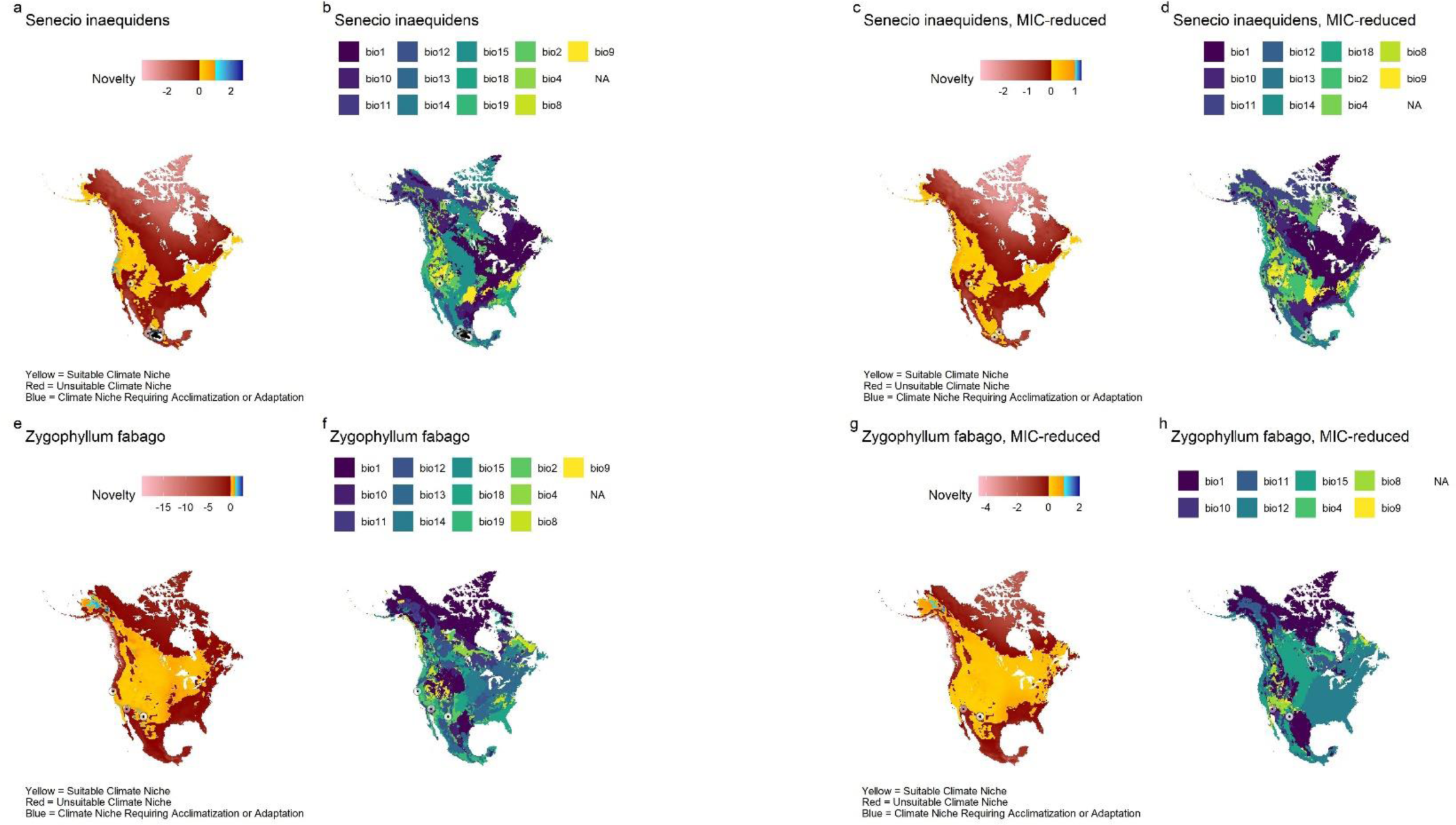
*– Model outputs for Senecio inaequidens* with the full variable set (a,b) and MIC-reduced set (c,d), and *Zygophyllum fabago* with the full variable set (e,f) and the MIC- reduced set (g,h). a, c, e, g) Climate suitability maps showing climate suitability in yellow, univariate climate suitability / type 2 novelty in blue, and unsuitable climate areas in red. b, d, f, h) Most important covariates contributing to model results. Black points surrounded by circles indicate observations of each species where the model estimates that the climate is unsuitable. Projection: NAS 1983 Equidistant Conic North America – ESRI:102010.

